# Size-scaling promotes senescence-like changes in proteome and organelle content

**DOI:** 10.1101/2021.08.05.455193

**Authors:** Ling Cheng, Jingyuan Chen, Yidi Kong, Ceryl Tan, Ran Kafri, Mikael Björklund

## Abstract

Senescent cells typically have an enlarged cell size but the reason for this has not been fully elucidated. As abnormal cell size may alter protein concentrations and cellular functionality, we used proteomic data from 59 unperturbed human cell lines to systematically characterize cell-size dependent changes in intracellular protein concentrations and organelle content. Increase in cell size leads to ubiquitous transcriptionally and post-transcriptionally regulated reorganization and dilution of the proteome. Many known senescence proteins display disproportionate size-scaling consistent with their altered expression in senescent cells, while lysosomes and the endoplasmic reticulum expand in larger cells contributing to the senescence phenotype. Analysis of organelle proteome expression identifies p53 and retinoblastoma pathways as mediators of size-scaling, consistent with their role in senescence. Taken together, cell size can alter cellular fitness and function through cumulative reorganization of the proteome and organelle content. An extreme consequence of this pervasive size-scaling appears to be senescence.

---

Maintenance of intracellular protein concentrations is important for cellular homeostasis at many levels. For example, cellular biochemical activities depend on the concentrations of metabolic enzymes, and dilution of specific cell cycle inhibitors may control cell size (*1–3*). Thus, changes in protein concentrations have the potential to dramatically alter normal cellular functionality. Organelle protein homeostasis may be particularly important as spatial compartmentalization of metabolic processes maintains optimal fitness and function under varying cellular demands (*4–7*) until proteostatic balance is gradually lost with aging (*8, 9*). Large cell size is one of the hallmarks of senescent cells (*10*), but it has been mostly considered as a passive outcome of the permanent cell cycle arrest which defines senescence (*17*). Excessive cell size may contribute to senescence (*12*), but how cell size or intracellular protein concentrations could be promoting senescence remains incompletely understood.

Intracellular concentrations of proteins are affected by synthesis and degradation rates but also by cell volume changes when cells grow or shrink (*2, 3, 13*). It is thought that the default behaviour for individual proteins and organelles is to increase proportionally with increasing cell size thus maintaining intracellular concentrations (Fig. 1A), whereas any protein displaying concentration changes could be a potential “cell size sensor” involved in the maintenance of cell size homeostasis (*3*). However, to what extent proteins or organelles obey or deviate from the constant scaling behaviour has not been systematically studied. We used publicly available data from the NCI60 cancer cell panel where the cell lines display an approximately tenfold range in volume (*14*) and for which a high-quality quantitative proteomic dataset is available (*15*) (Fig. 1B). Taking advantage of the natural size variation in these unperturbed, proliferating cell populations allowed us to avoid unwanted effects from drug-induced cell cycle arrest or cell sorting. The size variation in these 59 cell lines provides a larger dynamic range (Fig. 1B) than achievable in individual unperturbed cell lines. Furthermore, as the NCI60 cell lines originate from nine different tissues, any identified scaling behavior(s) should be largely independent of the cell and tissue type and exclude cell-type specific scaling.

**Fig 1.**
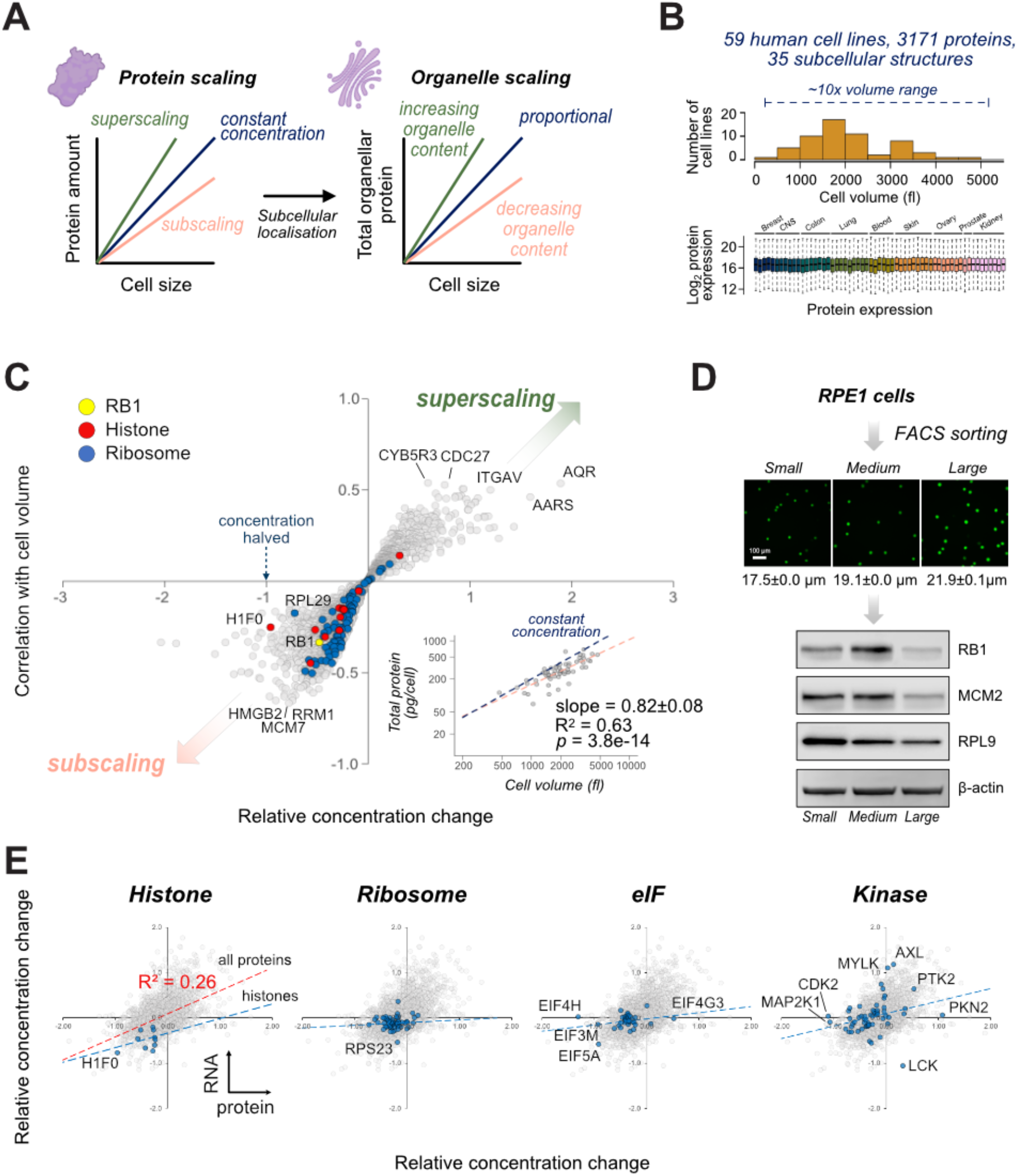
Scaling of the human proteome is ubiquitous and regulated primarily post-transcriptionally. **(A)** Schematic of protein and organelle scaling relations with cell size. **(B)** Cell volume and protein abundance distribution of the NCI60 cell panel. **(C)** Scaling of human proteome across the NCI60 panel. Each dot represents scaling of one of the 3171 detected proteins. Inset, total cellular protein measured by Bradford assay (*14*) as a function of cell volume in each of NCI60 cell lines. *p* value is for the linear regression model. **(D)** RPE1 cell sorting by size into small, medium and large fractions. An aliquot of cells was stained with Calcein AM tracer dye. Cell diameters of the subpopulations are indicated (mean±SD from 4 fields). For WB, 32 μg total protein was used for each lane. **(E)** Size-scaling of RNA (y-axis) vs. size-scaling of proteins, (x-axis). The grey dots represent the relative concentration changes (scaling slopes) for the whole proteome (3026 common RNAs/proteins, R^2^=0.26 in red), while histones, ribosome, eIF complex components and kinases are in blue. The regression model for all proteins is indicated with red dotted line, while the regression models for the indicated protein groups are in blue.

We first calculated changes in protein abundancies across the ten-fold range of cell volumes. The slope of the linear regression on the log-log scale provides the relative changes in protein concentrations, with the correlation coefficient describing how strongly the protein concentrations associate with cell volume changes (Fig. 1C, Table S1). This analysis revealed that cell size-related concentration changes are relatively small, but widespread. Nearly all proteins (3132/3171, 99%) displayed a relative change between −1 and 1, indicating that concentrations maximally halve or double when cells double in size as within a normal cell cycle. This scaling analysis confirmed the previously observed dilution of the retinoblastoma protein RB1 (*3*) as well as histones (*16*). Unexpectedly, these proteins were neither unique nor the most diluted ones. Many other proteins, including the ribosomal subunits were affected to similar extents (Fig. 1C). The proteins most diluted by scaling included High mobility group box 2 (HMGB2), ribonucleotide reductase catalytic subunit RRM1 as well as MCM7 and other minichromosome maintenance proteins (Fig. 1C, Table S1). The proteins whose concentrations increased with cell size included CDC27, a component of the anaphase-promoting complex, and CYB5R3, an antioxidant enzyme linked to longevity (*17*). The median relative concentration change for the 3171 proteins was −0.18; 95%CI = [−0.20, −0.17], indicating that most proteins are progressively diluted with increasing cell size. The overall proteome dilution could also be observed by plotting the previously measured total protein per cell (*14*) as a function of cell volume (Fig. 1C, inset). We further stained immortalized normal retinal pigmented epithelial (RPE1) cells for total protein using AlexaFluor 555-succinimidyl ester (*18*) and measured proteome scaling at the single cell level as a function of nuclear area as a proxy for cell size (*19, 20*). Proteome dilution in RPE1 cells occurred independently of changes in DNA content and yielded quantitatively similar dilution as the NCI60 proteome dilution (slope and standard error of 0.81±0.02, Fig. S1B). Consistent with the overall protein dilution, many protein complexes and other groups of proteins displayed dilution suggesting that such changes could have functional consequences (Fig. S1A).

To validate the proteomics data, we sorted RPE1 cells by flow cytometry into small medium and large subpopulations and analysed these by western blotting (Fig. 1D). Consistent with the proteomics data, we observed dilution of RB1, MCM2 as well as RPL9 as a representative of the ribosomal subunits within the ~2-fold average volume differences between the smallest and largest subpopulations. It has been reported that RB1 and histone concentrations are transcriptionally regulated (*3, 16*), but whether the overall proteome scaling is transcriptionally regulated is not known. To test this, we analysed scaling using NCI60 RNAseq data (*15*) and compared this to the proteome scaling. RB1 scaling is consistent at both RNA and protein level (slope = −0.4 RNA vs. −0.48 for protein) indicating transcriptional regulation. Histone scaling appears similarly transcriptional, in particular for the linker histone H1F0 (Fig. 1E). On the other hand, ribosomal proteins, eukaryotic translational initiation factor (eIF) subunits and the cell cycle regulatory kinases CDK2 and MAP2K1/MEK1 display dilution of the protein level despite maintaining constant RNA concentrations (Fig. 1E). Overall, RNA scaling explains only ~25% of the variance in proteome scaling suggesting that post-transcriptional regulation is dominant (Fig. S1, Table S2-S3). We further used gene ontology analysis and network analysis to identify strongly scaling cellular processes. DNA replication and cell cycle process were among the most enriched processes in the subscaling proteome (Fig. 2A, S1). Superscaling proteins include those involved with multicellular organismal process and extracellular matrix (ECM) organization, prolyl hydroxylation and lysosomal and endoplasmic reticulum proteins (Fig. 2B, S1). Of these, the cell cycle, prolyl hydroxylation, ion transport and nucleotide biosynthesis appear primarily post-transcriptionally regulated (Fig. S1).

**Fig 2.**
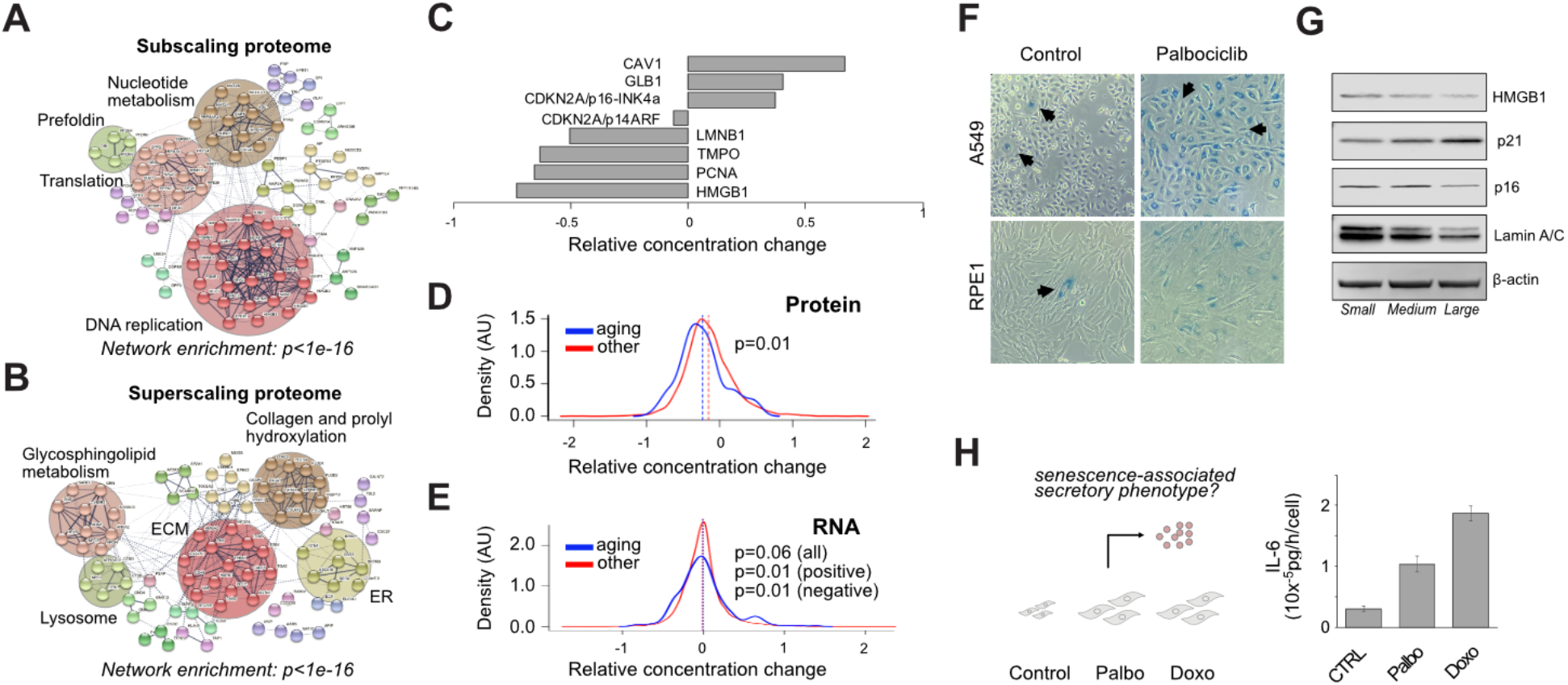
Senescence markers scale with cell size. **(A)** Connectivity analysis of subscaling proteome. Specific biological processes and protein groups are indicated. Network enrichment *p* value is compared to all genes as background set. **(B)** Connectivity analysis of superscaling proteome. **(C)** Scaling of selected senescence markers in proteomics data. **(D)** Scaling of known aging genes at protein level. **(E)** Scaling of known aging genes at RNA level. Significance (two-sided t test) for all, positively and negatively scaling aging genes compared to other genes is given. **(F)** Senescence-associated β-galactosidase assay for control and palbociclib treated A549 and RPE1 cells. Arrow indicate large senescent cells in control and small non-senescent cells in palbociclib culture. The cells were cultured for 6 days. **(G)** Western-blot analysis of senescence markers in RPE1 cells sorted by flow cytometry. Same lysates as in Fig. 1 were used, therefore β-actin blot is the same as in Fig 1D. **(H)** Schematic of senescence-associated secretory phenotype and analysis of secreted IL-6 amounts per cell in control, palbociclib and doxorubicin treated cells. Cells were incubated for four days, after which medium was changed and IL-6 secretion assayed after 8h by ELISA. Data shown is mean±SD, n=3.

What constitutes senescence remains incompletely defined as it can be induced by many distinct mechanisms (*10, 17*). Common hallmarks include large cell size, mitochondrial dysfunction, lysosomal accumulation, secretory phenotype as well as changes in plasma membrane composition and nuclear integrity, but what drives these changes remains incompletely understood. At the molecular level, a commonly used marker is senescence associated beta-galactosidase (SA-β-Gal) activity, but a persistent problem in understanding cellular aging is that no single marker is specific to senescence (*21*). We observed that size-scaling of many known senescence-related proteins was consistent with their accumulation or depletion in senescent cells. These included cell cycle markers (e.g., p16, PCNA), chromatin and nuclear lamina associated proteins (e.g., HMGB1, Lamin B1 and TMPO), plasma membrane caveolin (CAV1) as well as lysosome-associated GLB1, which is responsible for the SA-β-Gal activity (Fig. 2C, S2A). Cellular senescence and organismal aging are interlinked processes (*9*). The proteomics data included 98 high-confidence human aging-related proteins (*22*). These aging genes scaled significantly differently (p=0.01, two-sided t test) from the rest of the proteome becoming enriched into mostly subscaling portion, but some also in the superscaling part (Fig. 2D). At the RNA level, the 293 detected aging genes were significantly different from the rest of the RNA, although only when tested separately for sub- and superscaling RNAs (Fig. 2E).

As expected, SA-β-Gal activity was detectable in large RPE1 as well as A549 lung carcinoma cells incubated with CDK4/6 inhibitor palbociclib, but also in abnormally large cells in proliferating control cultures (Fig. 2F). However, SA-β-Gal activity can also be induced by other mechanisms than increased cells size. Palbociclib arrested RPE1 cells seeded at different densities showed highest numbers of senescent cells in densest cultures while having the smallest cells (Fig. S2B), perhaps due to lysosomal activation in response to nutrient limitation or contact inhibition.

To validate the size-scaling of senescence markers in proliferating cells, we further analysed the flow sorted RPE1 subpopulations by western blotting (Fig. 2G). Apart from p16^INK4a^, which is not induced in senescent RPE1 cells (*23*), the concentration changes were consistent with senescence (Fig. 2G). We next considered the senescence-associated secretory phenotype (SASP). SASP is a complex pro-inflammatory response, which can also affect neighbouring cells in a paracrine manner (*24*). Since the SASP response is hetereogenous in terms of the identity of the secreted proteins (*10*) and their intracellular levels are often low, many of the were not detected in all of the NCI60 cell lines. We therefore analysed IL-6, one of the most common SASP factors, in conditioned media from cultures with palbociclib- and doxorubicin-induced large cells. IL-6 secretion was significantly enhanced consistent with the senescence inducing ability of the large cell size (Fig. 2H).

Many organelles including mitochondria, lysosomes, and the nucleus display profound functional changes in senescent cells (*10*). Consistent with the idea that organelle changes can also be related to scaling, we have previously identified cell-size dependent changes in mitochondrial functionality (*6, 25, 26*). As organelles share a confined space, the analysis of organelles and other subcellular structures is conducive to understanding the functional consequences of scaling at the whole cell level (*27*). We therefore mapped the identified proteins in all cell lines to 35 subcellular structures based on Uniprot annotations and analysed the average expression of these subproteomes as a function of cell volume. For the majority of organelles, the slope on a log-log scale is close to one implying that the fraction occupied by most structures remain approximately constant (Fig. 3A, Table S4). Interestingly, the nuclear subproteome scales somewhat sublinearly (slope and standard error 0.94±0.02, p = 1.3e-54, R^2^=0.99), while the scaling of the nucleolus remains proportional to cell size (0.99±0.06, p =1.6e-24, R^2^=0.842)(Fig. S3A). This small deviation from isometry may explain why nuclear size “fails” to catch up with cell volume at extreme cell sizes (*28, 29*) as such a small difference becomes robustly observable only in very large cell sizes. The relative amounts of lysosomes, autophagosomes and endoplasmic reticulum (ER) increase superlinearly indicating expansion of these organelles in large cells (Fig. 3A, S3). Lysosomal scaling (slope and standard error 1.33±0.06) is quantitatively similar to the 1.3x scaling of yeast vacuole, the functional equivalent of mammalian lysosomes (*30*) suggesting evolutionary conservation.

**Fig 3.**
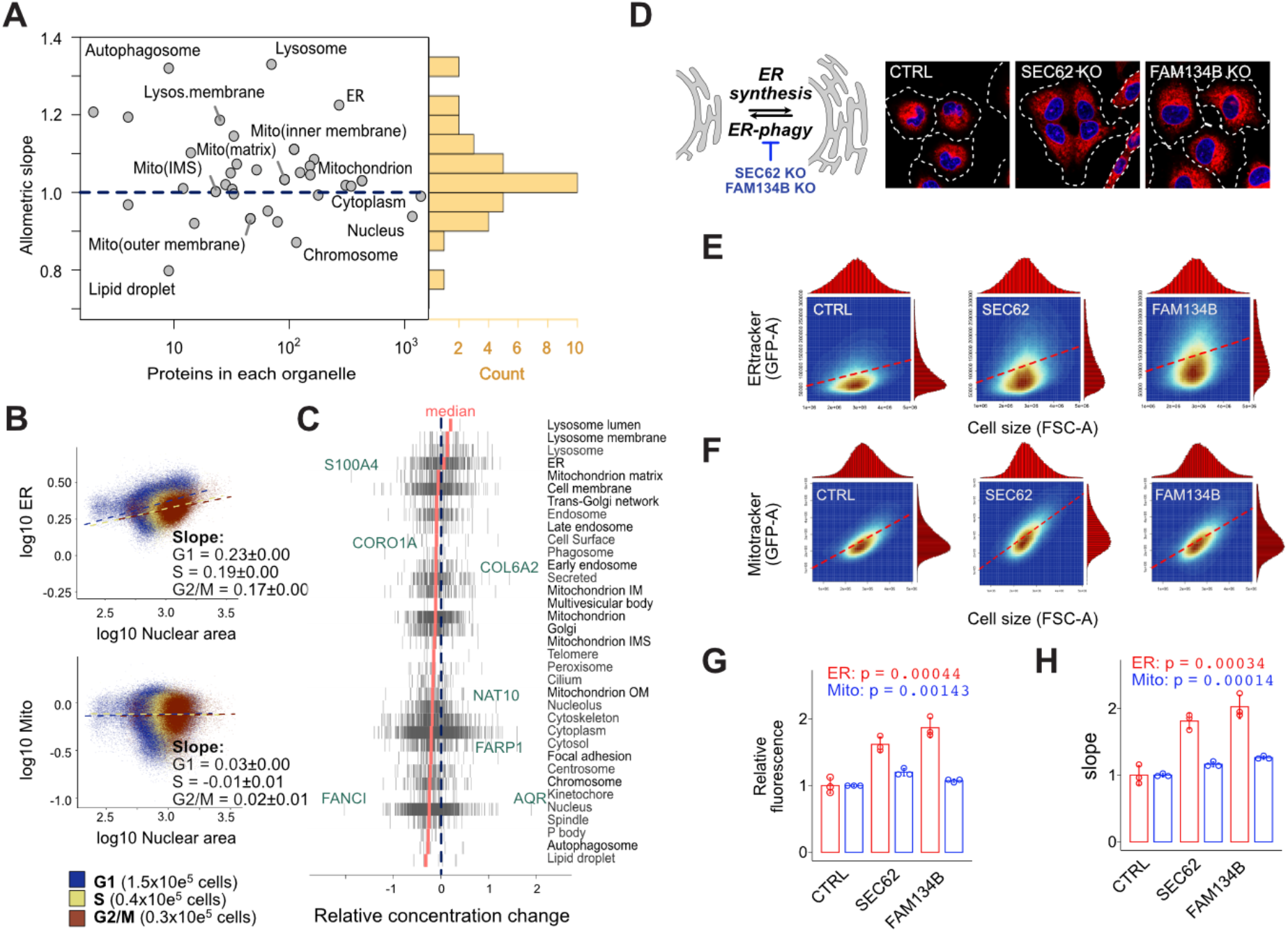
Organelle proteome scaling leads to senescence-like changes and ER expansion. **(A)** Scaling of 35 subproteomes using the NCI60 proteome data. The histogram depicts the distribution of the scaling exponents. For slope standard errors and summary statistics, see Table S4. **(B)** High-content microscopy analysis of normalised ER content (Concanavalin A-AlexaFluor488/ AlexaFluor555-SE) and mitochondrial content (Mitotracker Deep Red/AlexaFluor555-SE) scaling in RPE1 cells. Nuclear area was used as proxy for cell size. Blue, yellow and red colors indicate cell cycle phase for each cell based on DNA staining intensity. Regression models are shows for each cell cycle phase separately. In total ~2.2×10e5 cells were analysed. **(C)** Distribution of each protein (grey vertical lines) within organelle subproteomes. Selected differentially scaling proteins are indicated with green text. Median scaling of each subproteome is in red. **(D)** ER amount regulation by balanced synthesis and degradation by ER phagy. SEC62 and FAM134B knockout (KO) inhibit ER phagy leading to ER expansion (red = ER Tracker, blue = DAPI, white dotted line is cell outline) in A549 cells. **(E)** Flow cytometry analysis of ER scaling for control, SEC62 and FAM134 KOs. The red histograms show cell size and ER Tracker distribution. Red dotted line indicates linear regression. **(F)** Flow cytometry analysis of mitochondria scaling using Mitotracker green. **(G)** Quantification of relative fluorescence intensitity in SEC62 and FAM134B KO cells compared to control. **(H)** Relative change in scaling slope. Since the flow cytometry cannot directly measure cell volumes, scaling of control was normalised to 1. In G and H, *p* value is from ANOVA test. Data is mean±SD, n=3, each with ~1×10e5 cells.

The fraction occupied by ER is thought to remain constant (*4*). Consistent with the proteomics data, single cell analysis of ER content using concanavalin A staining in RPE1 cells also showed superscaling, which was largely independent of the cell cycle phase and consistent with the proteome scaling (Fig. 3B). Instead, scaling of mitochondria was isometric, consistent with the proteomic data and previous knowledge (*4, 25*). The superscaling of ER was further validated by comparing the scaling of the common ER marker SEC61B with the lysosomal marker LAMP1 (slopes in proteome data = 0.41 and 0.30, respectively) endogenously tagged with mNeonGreen in A549 cells (Fig. S3E). Flow cytometry confirmed their superscaling relative to the total protein (Fig. S3F). To find evidence for ER scaling *in vivo,* we searched for literature where cell and organelle size has been systematically measured in a tissue context. In chemically induced carcinogenesis models in rat livers, preneoplastic foci display increased cell size compared to the non-focal unaffected areas (*31, 32*). Reanalysis of this data showed that the fractional volume occupied by the nucleus is reduced in large cells in the focal areas compared to the smaller cells in non-focal tissue, but the logarithm of the nuclear and cell volume remained constant suggesting scaling (Fig. S4A-C). Among the measured organelles, the volume occupied by ER increased most substantially, from 21% to 32% in large cells (Fig. S4D).

Proteomic data also allows the identification of key differences in organelle composition, which is not easily achievable by microscopy approaches. The scaling of individual proteins occasionally varies quite substantially (Fig. 3C) suggesting cell size-dependent differences in organelle composition even when their overall scaling is proportional to cell size. Such changes may potentially be important for organelle function. An example is the nucleolus, which contains strongly superscaling NAT10 (Fig. 3C), an N-acetyltransferase associated with premature-aging in Hutchinson-Gilford progeria syndrome (*33*). Overall, our analysis of subcellular organelle proteome scaling recapitulates known scaling relations as well as identifies ER as a superlinearly scaling organelle.

A simple mechanism for achieving superlinear scaling of ER would be to alter the balance between ER synthesis and degradation by ER-phagy (Fig. 3D)(*34–37*). To test this, we knocked-out two of the key regulators SEC62 and FAM134B (*35–37*) and measured scaling of ER and mitochondria using flow cytometry (Fig. 3E, F). Individual ER proteins displayed increased expression in the KO cells (Fig. 3G,H) and both knockout cell lines resulted in an approximately 1.5 to 2-fold increase in ER tracker, but not in Mitotracker signal, as well as a larger slope, indicating altered scaling (Fig. 3G, H). The superlinear scaling of ER is consistent with the symmorphosis hypothesis, which predicts that a part of a system should quantitatively match the overall functional demand (*38*), in this case the secretory phenotype.

Detailed analyses of individual proteins can allow the identification of mechanisms that contribute to scaling, but selecting the most important candidates for testing is not trivial. Furthermore, as individual organelles are embedded in the context of the whole cell, their expansion or contraction can affect the scaling of other structures. To begin unraveling this, we first visualised the inter-relationships of all the 35 subcellular structures, which revealed that the cell surface is the most distinct subproteome displaying the poorest correlations with other cellular structures (Fig. S5A). Comparison of linear and allometric models suggested that unlike other subcellular proteomes, the variance of the errors in cell surface proteome scaling are a better fit with linear than allometric power-law (Fig. S5), while speculative this could be related the more flattened shape of the large adherent cells, which is another morphological characteristic of senescence (*10*).

The ability to reduce complex cellular morphometric shape space into a few principal components (*4, 39*) suggests that the major mechanisms of how cell size affects proteome reorganization may be less complex than anticipated. We next sought statistical evidence for this by associating subcellular proteome levels with the essentiality of individual genes from systematic genome-wide CRISPR-Cas9 knockout screens of 263 cell lines for which proteome data is available (Fig. 4A, Table S5) (*40, 41*). Clustering all the organelle associations with ~18 000 gene knock-outs using Uniform Manifold Approximation and Projection (UMAP) yielded the Organelle Scaling Dependency Map (Fig. 4B). This map illustrates the similarities and differences in genetic dependencies for the subcellular structures. For example, the nucleus and the lysosome display inverse dependencies (Pearson correlation R =-0.43, Fig. 4B) as is expected from the opposite scaling behaviour of these organelles (see Fig. 3A). As another example, the nucleolus clusters and correlates more closely with mitochondria (R=0.59) than the nucleus with mitochondria (R=0.21)(Fig. 4B) suggesting coupling of energy metabolism and ribosome biogenesis for protein synthesis and growth. Analysis of the genetic dependencies identified that nuclear and lysosomal protein levels associate strongly with p53 and retinoblastoma pathways, the two most understood senescence regulators (*11*). In addition to p53 (TP53), we observed p53 binding protein (TP53BP1), p53 deubiquitinase USP28 and DNA damage sensor ATM kinase as well as retinoblastoma pathway components RB1 and p21 (CDKN1A) among the top dependencies. MDM2, the E3 ligase and negative regulator of p53 as well as Cyclin D1 (CCND1) KO displayed opposite associations compared to p53 (Fig. 4B, S6A). Among all subcellular structures, early endosome subproteome levels displayed the strongest dependencies on p53 (Fig. S6B). Reduced SA-β-gal staining in USP28 knockout cells (Fig. 4C) is consistent with our association analysis and with previous identification of USP28 as a senescence regulator (*42*).

**Fig 4.**
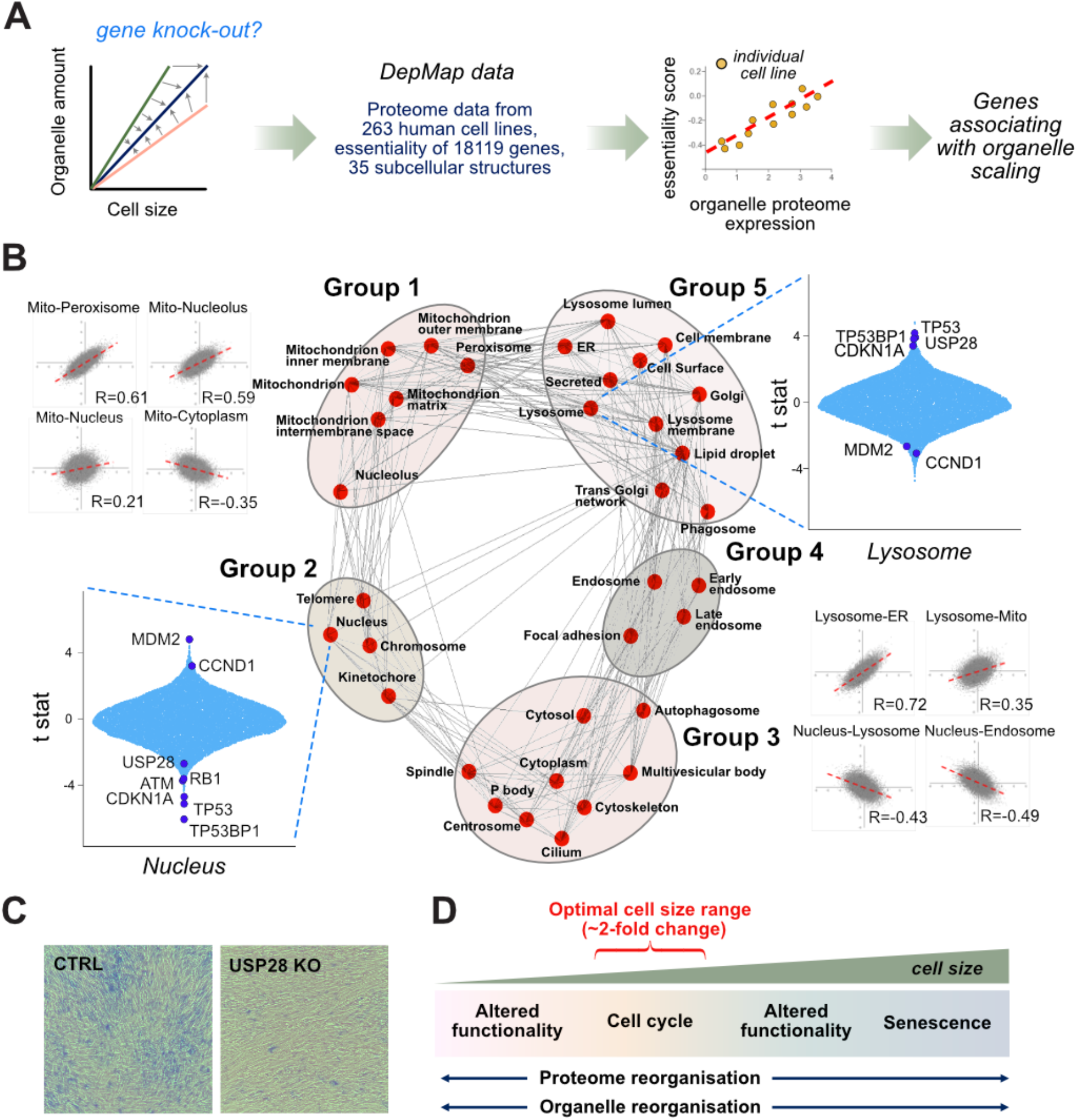
Organelle Scaling Dependency Map identifies p53 and RB pathways as contributors to scaling. **(A)** Schematic of the organelle subproteome scaling analysis based on DepMap gene essentiality and proteomics data from 263 cell lines. **(B)** The central network is the Uniform Manifold Approximation and Projection (UMAP), which divides the 35 subproteomes into five subgroups. The genetic dependencies for each of the ~18000 gene knockouts (gray dots) for selected individual pairs of subproteomes are shown along with their Pearson correlations. The violin plots display the genetic associations of each of the knockouts. The top scoring retinoblastoma and p53 pathway components are indicated with dark blue. **(C)** SA-β-gal assay of RPE1 control and USP28 KO cells. Cells were incubated for 9 days before fixation and staining **(D)** A model for the consequences of cell size scaling.

Interestingly, the transcription factors involved in lysosomal and mitochondrial biogenesis (*43, 44*) appear to be less important determinants for organelle amount that p53 and RB pathway components (Fig. S6C). For example, lysosomal protein levels were hardly associated with TFEB and TFE3, possibly due to functional redundancy. Similarly, mitochondrial proteome levels displayed only a weak dependency on the main mitochondrial biogenesis factor PGC1-alpha (PPARGC1 KO). Indeed, mitochondrial and lysosomal proteomes were more dependent on many individual PI3K-AKT-TOR pathway components than their biogenesis factors (Fig. S6C). Altogether, the Organelle Scaling Dependency Map provides statistical association of proteome scaling with p53 and retinoblastoma pathways across hundreds of cell lines, consistent with their role in senescence, but many additional proteins are potentially involved in scaling (Fig. S6D, Table S5).

We have studied the scaling of the human proteome using data from tens of unperturbed, normally proliferating cell lines. Size-scaling appears to be a systems-level attribute, which is regulated at multiple levels, although the dilution of ribosomes may be a major culprit for post-transcriptional regulation. Our analysis highlights the pervasive nature of scaling at the level of individual proteins as well as organelles and implies that cell size has important consequences on optimal cellular functionality (*25*). For most of the proteins, cell-size dependent concentration changes are relatively small, allowing cells to grow and divide within their optimal range of cell sizes without adverse functional effects from protein scaling. Because many of the most strongly scaling proteins are related to cell cycle and DNA replication, scaling could be beneficial within the two-fold volume range of the normal cell cycle (*1–3, 45*) whereas the antagonistic effects, including senescence, would manifest beyond a much larger cell size range (Fig. 4D).

While large cell size has long been recognized as one of the central hallmarks of senescent cells, the relevance of cell size on the senescence program has been overlooked. Our observations suggest that enlarged cell size may induce senescence. Together with similar findings reported by the Skotheim group (*16*), our data is consistent with the hypothesis that cell size promotes a senescence-like program through size-scaling even in the absence of a cell cycle arrest. This may in part explain the long-standing paradigm of why there are no specific markers for senescence (*10, 11, 21*). Overall, these observations challenge some our current understanding on cellular aging and prompts us to reassess some of the underlying assumptions of senescence. Critically, our observations suggest that cell size not only impacts the functionality of the abnormally large cells themselves, but potentially also the neighbouring cells through the cell-size/senescence-associated secretory phenotype. Therefore, being of the right cell size is likely to be important for the fitness of the whole organism.

## Acknowledgments

We thank Tiannan Guo for NCI60 proteome data, Philipp Kaldis and Teemu Miettinen for critical comments, Zongsheng Jiang for help in CRISPR cell line generation and Chan Kuan Yoow for USP28 KO cells.

## Author contributions

Conceptualization: M.B.; Methodology: C.T., R.K. and M.B.; Investigation:L.C., J.C, Y.K, C.T, M.B, Writing – original draft: M.B.; Writing – reviewing and editing: all authors; Visualization: J.C. and M.B

## Data and materials availability

The underlying proteomics and RNAseq data are publicly available from https://discover.nci.nih.gov/cellminer/home.do or ref 10, and https://depmap.org/portal/. All other data are available as supplementary materials.

## List of Supplementary materials

Materials and Methods

Figs. S1 to S6

Tables S1 to S5

## Supplemental Information

### Methods

#### Proteome scaling analysis

NCI60 cell line diameters derived from image based analysis (*1*) and proteomic dataset (*2*) was processed as follows; we estimated cell volumes by assuming that cells are approximately spherical after trypsin treatment. Therefore, cell volume is V=4/3π(d/2)3, where d is the average reported diameter. To obtain protein concentrations for each of the 3171 proteins across 59 cell lines, the total signal intensity from mass spectrometry was normalized first to the average protein signal across the whole dataset and further to total cellular protein measured separately for each cell line (*1, 3*). All available data from the 59 cell lines was included in the analysis. The resulting protein abundancies were divided with the cell volumes to obtain concentrations for individual proteins. To obtain organelle protein concentrations, we downloaded annotated localisations from UniProt database (https://www.uniprot.org/locations/), including 35 common organelles and other subcellular structures. We included only reviewed localizations using the following type of SPARQL query locations:(location: “Lysosome [SL-0158]”) AND reviewed:yes AND organism:“Homo sapiens (Human) [9606]”. The protein abundancies for each protein at each organelle were summed and used as total organelle subproteome.

Regression analyses were performed using RStudio 1.4.1106 (R base version 4.1). For allometric analysis regression analyses were done using *lm* linear model for log10 transformed data. Log10 values of cell volumes and protein concentrations were used to obtain relative concentration changes. In this way, proportional scaling with cell volume yields a slope of zero.. Model statistics, including slope, slope standard error and R2 values were extracted from the *lm* summary output. For organelle analysis, we used typical allometric analysis where organelle mean protein abundancies rather than concentrations were used. In this case, the proportional scaling yields a value of one. To test that allometric power law-based analysis is a valid description for organelle scaling we compared it with linear model on non-logaritmic scale. Model evaluation included comparison of *p* value, R^2^ and Akaike information criterion (AIC, smaller AIC indicates a better fit). We also considered one of the main assumptions behind regression analysis, constant variance of the residuals (difference between fitted model and measurement). To evaluate linear model validity, statistical testing of residuals’ variance was done using Breusch-Pagan variance tests in the *lmtest* package (v. 0.9-37).

Rug plots for individual proteins within an organelle or protein complex were plotted using *geom_rug* function in *ggplot2* package. The scaling of each individual protein is indicated with grey vertical line. Median scaling was calculated and included in the plot as red solid line.

#### RNA scaling analysis

The processed RNAseq data for NCI60 cells (*2*) was used without normalisation to the total cellular RNA. For RNAseq data, log2 transformed cell volumes were used as the original RNAseq data was expressed on a log2 scale. Otherwise, the scaling analysis was done similarly to the protein analysis. The protein and RNA datasets were combined resulting in 3026 genes with both RNA and protein scaling data.

#### Gene Ontology and other common bioinformatics analyses

Gene Ontology analysis was performed using GOrilla (*4*). Data was plotted using either *ggplot2* package (v. 3.3.2) in R, GraphPad Prism 8 or Microsoft Excel. Network analysis was performed using String database (version 11.0)(*5*).

#### DepMap association analysis

Gene essentiality scores and DepMap proteome data were downloaded from Broad Institute’s DepMap portal (https://depmap.org/portal/download/). The DepMap Public 20Q2 version were used for Gene essentiality scores (*6*). This data contains copy-number corrected, scaled and normalized results of genome-scale CRISPR knockout screens expressed as gene-level CERES scores for 18119 genes in 769 cell lines, of which 236 also have proteome data containing expression levels of 5153 proteins (*7, 8*). All available datapoints were included in the analysis. The proteome data was summarized for each of the 35 organelles as described for the NCI60 proteome data. The associations were expressed using the T statistic, which retains the directionality of the association. For visualization of the organelle proteome level dependency associations, Uniform manifold approximation and projection (UMAP) was used. Briefly, data was imported into Spyder 4.2.0 environment (based on Python 3.7). UMAP algorithm was run using *umap* package (v. 0.1.1). The default mapping parameters were used with *n_neighbors=15* and *min_dist=0.01.* The *n_neighbors* determine the size of neighboring sample points, while *min_dist* is the minimum distance among points in each cluster. To identify the individual organelles in the UMAP, *umap.plot.interactive* function was used while the connectivity between the organelles was plotted using *umap.plot.connectivity* function with default settings.

#### Cell culture

RPE-1, HL-60 and A549 cells were obtained directly from ATCC. A549 and HL-60 cells were cultured in 5% FBS in RPMI1640 (GE Life Sciences, SH30096.01) supplemented with 1mM glutamine. hTERT-RPE1 cells were cultured in 10% FBS in DMEM containing 25 mM glucose (Biological Industries, 06-1055-57) or for the high content microscopy experiments in 10% FBS in DMEM/F12 medium (Life Technologies). All cell lines were cultured in the presence of penicillin and streptomycin and tested to be negative for mycoplasma.

#### CRISPR knockout and knock-in

sgRNAs targeting SEC62 were cloned using (F: CACCGagaaaaacctgatcatcatg and R: AAACcatgatgatcaggtttttctC) and for FAM134B using (F: CACCGAACACGCCCAAGTATCATGA and R: AAACtcatgatacttgggcgtgttC). These were cloned into BsmBI-v2 digested plentiCRISPR V2 (addgene, #52961). Non-targeting control sgRNA (GGGCGAGGAGCTGTTCACCG) was purchased from Genscript. The resulting plasmids were transfected into A549 cells with jetPRIME (Polyplus transfection, #114-15) and selected with 0.75ug/ml Puromycin dihydrochloride hydrate (BBI, A610593-0025). Single-cell clones were obtained by limiting dilution. USP28 KO cells were a gift from Dr. Chan Kuan Yoow.

For mNeonGreen as C terminal fusion with SEC61B and LAMP1, knock-in cassette consisted of the left homologous arm-mNeonGreen-T2A-puroR or G418 resistance gene-right homologous arm was cloned in pUC19 vector. SEC61B knock-in cells were selected with 0.75ug/ml Puromycin and 1mg/ml Geneticin (G418 Sulfate, Selleckchem, S3028). LAMP1 knock-in cells were selected with 0.75ug/ml Puromycin only. The sgRNA sequences used for SEC61B were CTCCTTCTTCAGATGACAGA and atccatctgtcatctgaaga. Left homo arm was amplified with F:gaccatgattacgccaagcttTGTAGCCCATTGCTCTTCAGGG and R:tcctccCGAACGAGTGTACTTGCCCC and the right homo arm with F:tcttctgaTGAAGAAGGAGGAAAAAACCCAAC and R:aaaacgacggccagtgaattcAGGGCGTACCAGAAACTGAC. The sgRNA sequence used for LAMP1 was CTATCTAGCCTGGTGCACGC. Left homo arm was amplified with F:gaccatgattacgccaagcttGACGCCAGAGGTGAGAACCC and R:tcctcctccGATAGTCTGGTAGCCTGCGTGAC and the right homo arm with F:tgcctgaCCTGGTGCACGCAGGCAC and R:aaaacgacggccagtgaattcACACCCTTCTCTCTGCACCGAG. The sgRNA and knock-in cassette plasmids were co-transfected into A549 cells with jetPRIME.

#### Measurement of cell size

For a typical experiment on a 6 well plate, 0.5-1.0×10^5^cells/ml were plated in 2 ml complete culture medium containing 50 nM doxorubicin or 1 μM palbociclib and cultured for the time indicated. To measure cell size, cells were trypsinized with 200 μl TrypLE Express (Thermo-Fisher) and mixed 1:1 with 0.2% Trypan Blue in PBS as viability dye. Cell number, diameter and viability measurements were obtained using Countstar BioTech Automated Cell Counter (ALIT Life Science, Shanghai, China).

#### Flow cytometry and cell sorting

Proliferating HL-60 cells were washed with PBS, resuspended in staining solution containing 100 nM Mitotracker Green (CST, 9074P) or 500 nM LysoBeacon Green (Genecopoeia, C048) for 15 min at 37°C together with 0.5μM CellTracker Deep Red (ThermoFisher, C34565) as amine-reactive dye to label total cellular protein. Cells were washed once with PBS and analysed by ACEA Novocyte flow cytometer. Typically, 1-2×10^5^ cells were analysed per sample. Scaling analysis the function of cell size (FSC-A) was done as previously described (*9*). Briefly, single-cell fluorescence ratio of green (measured scaling variable) and farred signal (total proteome) was binned by 50000 FSC-A units and median values for each bin calculated and plotted (dots). Locally weighted regression analysis was performed with the *loess* function in R and the regression (solid line) plotted with 95% confidence interval (shaded line). The cell size distribution by FSC-A is shown for reference. For scaling of SEC61B and LAMP1 knock-in cells, the cells were directly stained with CellTracker Deep Red and processed as above. For experiments involving A549 control, SEC62 and FAM134B cells, the cells were stained for 30 min with 1μM ER tracker green (BODIPY™ FL Glibenclamide, Beyotime, C1042S) or 400 nM Mitotracker Green together with CellTracker Deep Red. The cells were trypsinized, collected by centrifugation and processed as above.

For sorting of RPE1 cells, BD Influx Cell Sorter was used. RPE1 cells were trypsinized and resuspended in PBS with 1 mM EGTA. Preliminary sorting experiments showed that this instrument separates cells by size better using side-scatter SSC than forward scatter FSC. The cells were sorted into three populations including lowest, middle and highest 15% of the SSC scatter signal of the 488 nm laser. We sorted ~1 million cells per fraction and collected them by centrifugation at 800xg. An aliquot of each fraction was used to measure average cell diameter with Countstar Cell Counter or stained with 1μm Calcein FM (Beyotime, Shanghai, China) for 10 min and imaged with 20x magnification using Etaluma LS620 fluorescence microscope. The remaining cells were lysed and analysed by western blotting as detailed below.

#### Western blotting

Cells were washed with PBS and lysed with 1% NP40-PBS for WB. Proteins were separated on 4-12% or 4-20% SurePAGE Bis-Tris gels (Genscript) and transferred on PVDF membrane (Merck Millipore, ISEQ00010). Membranes were blocked with QuickBlock™ Blocking Buffer (Beyotime, P0252), incubated with primary antibodies in QuickBlock™ Primary Antibody Dilution Buffer (Beyotime, P0256), followed by HRP (CST, #7074) or AlexaFluor Plus 680 or 800 conjugated secondary antibodies (Invitrogen, A32735 and A32729). The signals were detected with LiCOR Odyssey imager.

The following antibodies were used: RPL9 antibody [EP13752] (Abcam, ab182556), MCM2 antibody [EPR4120] (Abcam, ab108935), Rb (D20) Rabbit mAb (CST, #9313), HMGB1 (D3E5) Rabbit mAb (CST, #6893), p16 INK4A (D3W8G) Rabbit mAb (CST, #92803), p21 Waf1/Cip1 (12D1) Rabbit mAb (CST, #2947), Lamin A/C Rabbit mAb (ABbclonal, Wuhan, China, A19524), Calreticulin (D3E6) XP Rabbit mAb (CST, #12238), Calnexin (C5C9) Rabbit mAb (CST, #2679), ERp72 (D70D12) XP Rabbit mAb (CST, #5033), Anti-SEC62 antibody [EPR9212] (Abcam, ab137022), beta-Actin mouse mAb (Genscript, A00702).

#### Senescence assays

Senescence associated beta-galactosidase (SA-β-Gal) activity was performed by a commercial kit (Beyotime, C0602). Cells were cultured in complete medium for the time indicated with 1 μM palbociclib or 50 nM doxorubicin in 6 well plates before fixation and staining. For confluency dependent senescence, cells were plated between 0.25e5 and 1.5e5 cell/ml. For IL-6 assay, RPE1 cells were cultured for 3 days, fresh medium changed and IL-6 secretion measured after 8h. Conditioned medium was diluted 1:4 and analysed by human IL-6 ELISA kit (ABclonal, RK00004).

#### Live cell confocal microscopy

For SEC62 and FAM134B knockout cells, ER-Tracker Green (Beyotime,C1042S) was diluted to 1 μM working concentration together with 1:1000 dilution of Hoechst 33342 (Beyotime,C1027) in ER-Tracker Green dilution buffer (Beyotime,C1042S). Cells were incubated with 1ml staining solution for 30min at 37C°, 5% CO2. Cells were washed twice with culture medium and imaged using Zeiss LSM880 confocal microscope. The SEC61B and LAMP1 mNeonGreen knock-in cells were stained with Hoechst 33342 as above.

#### High content microscopy

RPE1 cells were seeded at 3000 cells/96-well (PerkinElmer CellCarrier-96 Ultra Microplates) and incubated for 24h. 30 min before fixation, cells were incubated with 375nM Mitotracker Deep Red (Invitrogen, M22426) at 37°C, 5% CO2. Cells were fixed in 4% paraformaldehyde (Electron Microscopy Sciences, Hatfield, PA) for 10 min, followed by permeabilization using 0.1% Triton X-100 dissolved in HBSS for 15 min at RT. ER was stained with 100μg/mL concanavalin A (ConA) conjugated to AlexaFluor 488 (Invitrogen, C11252) in 1% BSA/HBSS for 30 min at RT. Cells were then stained with 0.4 μg/mL AlexaFluor 555 carboxylic acid, succinimidyl ester (Invitrogen, A-20006) for 2 hr at RT. DNA was stained with 1 μg/mL DAPI (Sigma, D8417) for 10 min at RT. Cells were imaged using the Operetta High-Content Imaging System (PerkinElmer, Woodbridge, ON) at 20X magnification. Image processing was performed using custom-written tools in MATLAB. DNA, ER and mitochondrial content were determined by quantifying the integrated intensity of DAPI, ConA and MitoTracker, respectively, on a single cell basis.

**Fig. S1.**
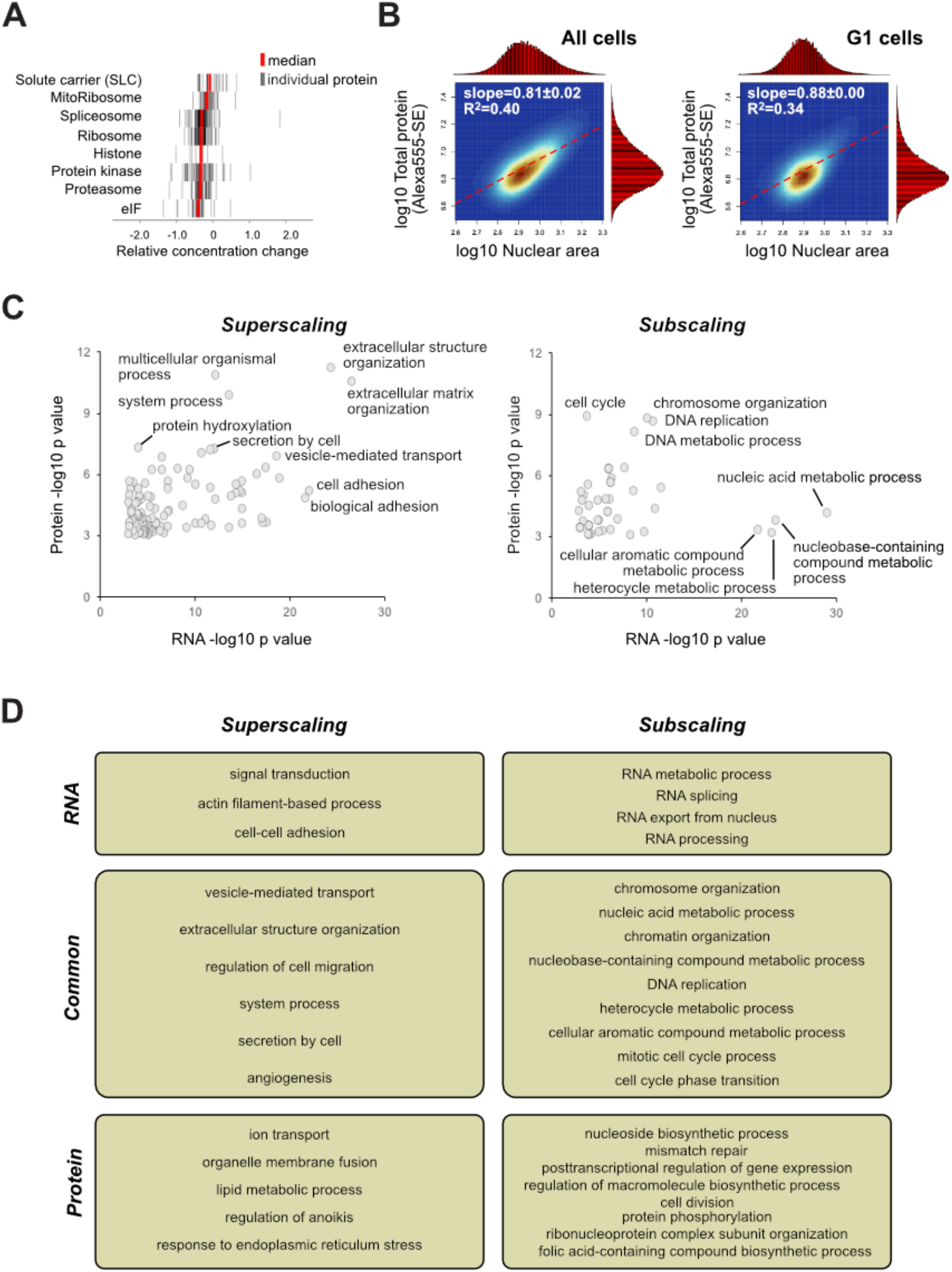
Human proteome dilution and regulation scaling. **(A)** Rug plots showing scaling of individual proteins (grey bars) and median (red) for each group of proteins and protein complexes. (B) Dilution of RPE1 proteome (left panel = all cells, right = G1 cells) analysed by amine-reactive AlexaFluor 555-succinimidyl ester staining using high-content microscopy. The red histograms show cell size and ER Tracker distribution. DAPI staining was used to quantify nuclear area and classify the cell cycle phase based on DAPI intensity. Approximately 2×10e5 cells were analysed. **(C)** Gene ontology comparison of superscaling and subscaling at protein and RNA level. **(D)** Enrichment of gene ontology groups which are enriched in RNA only (but no significant scaling at proteome level), commonly enriched in both RNA and protein data (transcriptionally regulated scaling) and enriched in protein level only (post-transcriptionally regulated scaling).

**Fig. S2.**
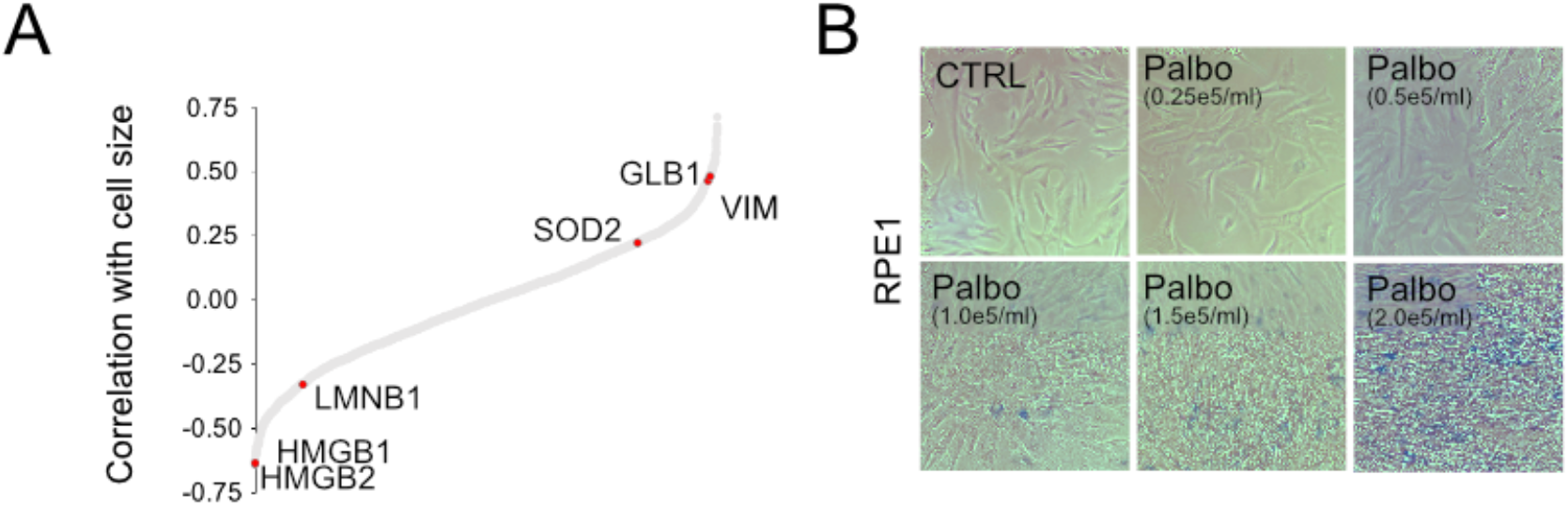
Senescence markers are altered by cell size. **(A)** Cell size correlations (Pearson R) for the proteins detected in the NCI60 proteome dataset. Selected senescence associated proteins are indicated in red. **(B)** Cell density dependent induction of senescence in RPE1 cells treated with 1 μM Palbociclib. Cells were stained after 4 days for SA-β-Gal.

**Fig. S3.**
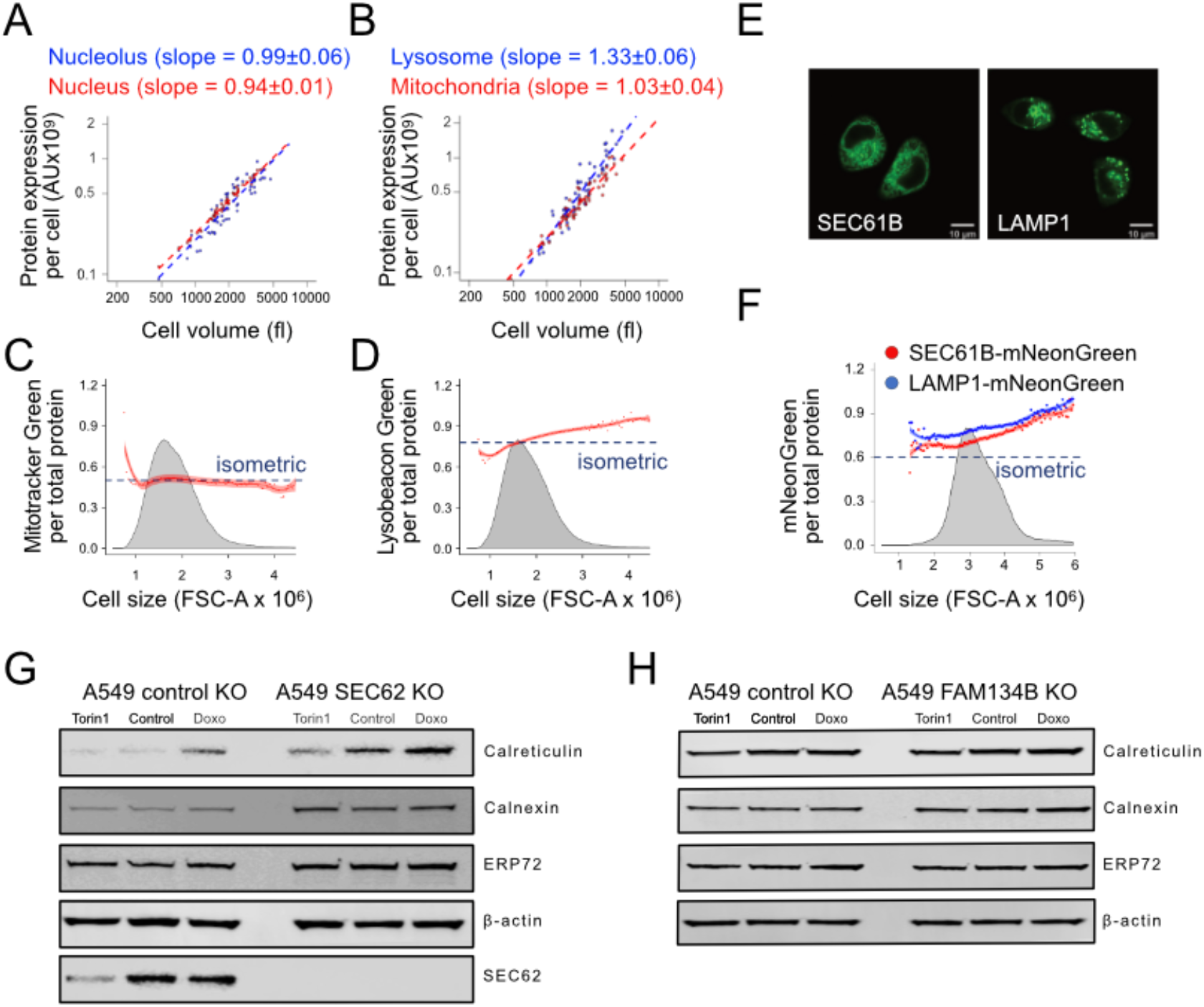
Validation of organelle scaling. **(A)** Scaling of nucleolar (blue) and nuclear (red) proteomes in NCI60 data. Each dot represents one of the NCI60 cell lines, the linear regressions are shown. The data shows slopes and standard error. **(B)** Scaling of lysosomal (blue) and mitochondrial (red) proteomes in NCI60 data. **(C)** Mitotracker green scaling in HL60 cells by flow cytometry. The cell size distribution of the population is in grey, the Mitotracker signal normalized to amine-reactive Celltracker Deep Red dye at single cell level is in red. Isometric scaling is shown for reference as blue dotted line. For detailed description of the analysis, see (*9*). **(D)** Same as panel C, but cells were stained with Lysobeacon green lysosomal dye. Approximately 1e10^5^ cells were analysed. **(E)** Confocal images of SEC61B and LAMP1 knockin with mNeonGreen in A549 cells. Scale bars are 10 μm. **(F)** Scaling of SEC61B and LAMP1 knock-in cells analysed by flow cytometry with normalization to Celltracker Deep Red dye signal. For panels C, D and F, approximately 1e10^5^ cells were analysed. **(G)** Scaling of ER resident proteins Calreticulin, Calnexin and ERP72 in A549 control and SEC62 knockout cells. Cells were treated for 3 days with 100 nM Torin-1 or 10μM palbociclib to decrease and increase cell size, respectively. b-actin was used to control for equal protein loading. **(H)** Scaling of ER resident proteins in A549 control and FAM134B knockout cells. No functional antibody for FAM134B antibody was available so knockout was validated by DNA sequencing (data not shown).

**Fig. 4.**
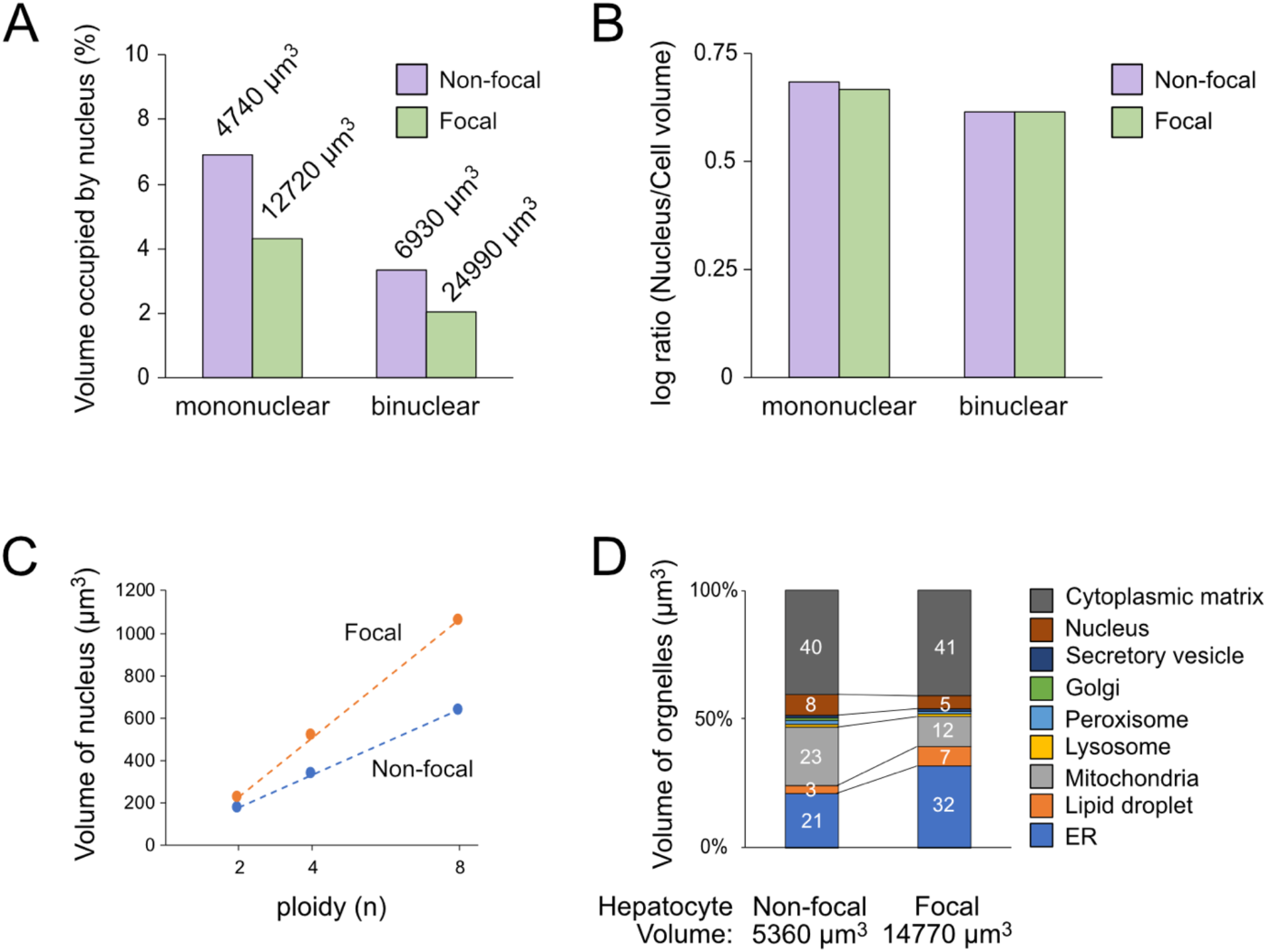
ER superscaling *in vivo.* **(A)** Volume fraction of nucleus in mononuclear and binuclear rat hepatocytes in adjacent normal (non-focal) and pre-neoplastic (focal) lesions. Average cell sizes are indicated above the bars. **(B)** Constant nuclear to cell volume ratio indicating proportional scaling. **(C)** Scaling of nuclear volumes in focal and non-focal cells differing by ploidy. **(D)** Volume fractions of various organelles in non-focal and focal hepatocytes. The data was reanalysed and plotted from ref. (*10, 11*).

**Fig. S5.**
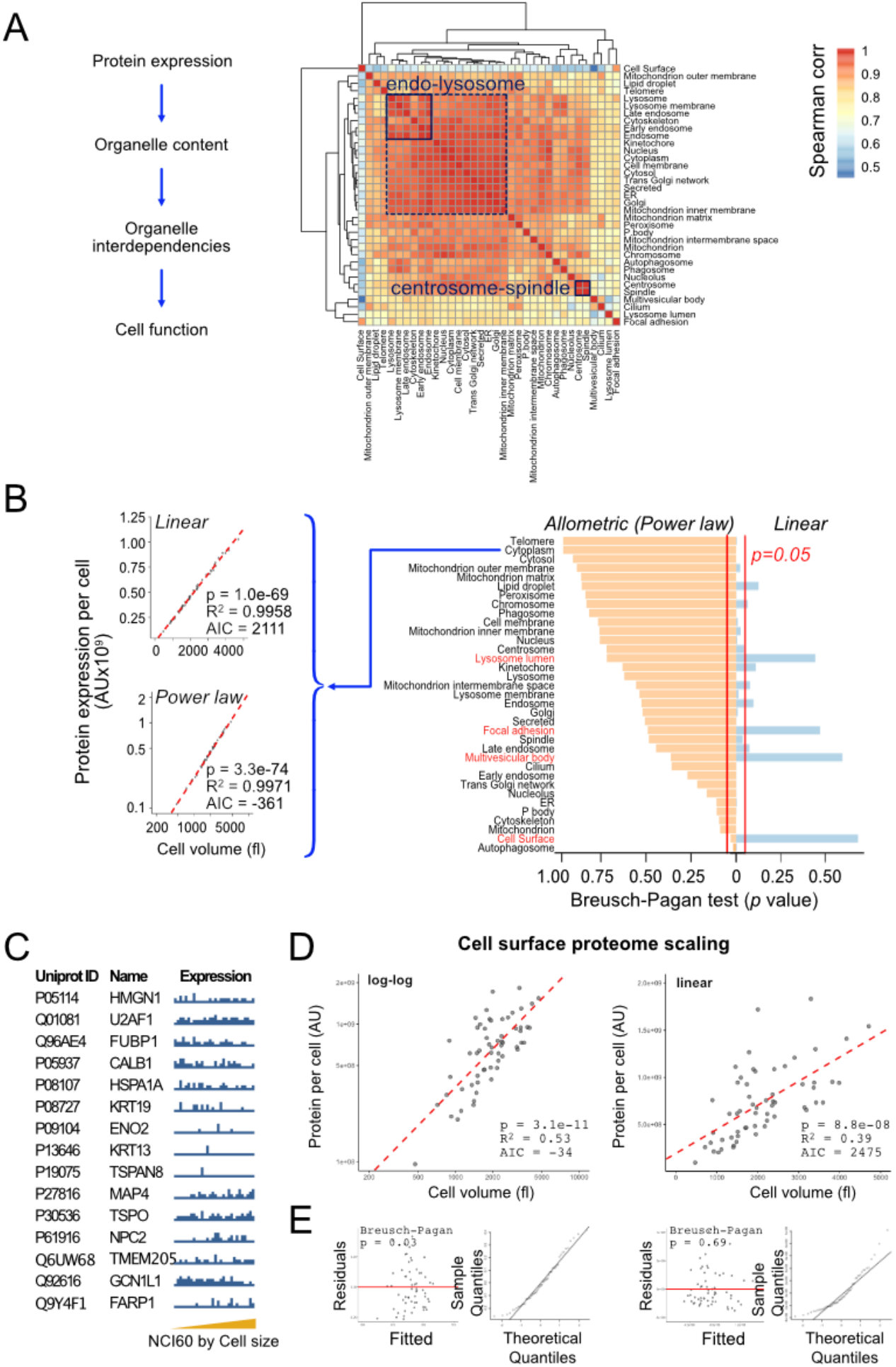
Analysis of organelle interdependencies and identification of cell surface as a differentially scaling subproteome. **(A)** Spearman rank correlations matrix of the median expression levels of each subproteomes across the NCI60 cell lines. **(B)** Comparison of scaling models on linear and allometric/log-log power law scale using Breusch-Pagan test for heteroscedasticity of the residuals. Subproteomes that fit linear scaling model are indicated in red. The left panels show cytoplasmic subproteome scaling and corresponding summary statistics. **(C)** Expression of annotated cell surface proteins across the NCI60 cell lines sorted by cell size. **(D)** Comparison of the allometric log-log and linear scaling for cell surface subproteome. The regression models for are shown with red dotted line along the summary statistics. **(E)** analysis of the residual fits and quantiles for the allometric log-log and linears in panel D.

**Fig. S6.**
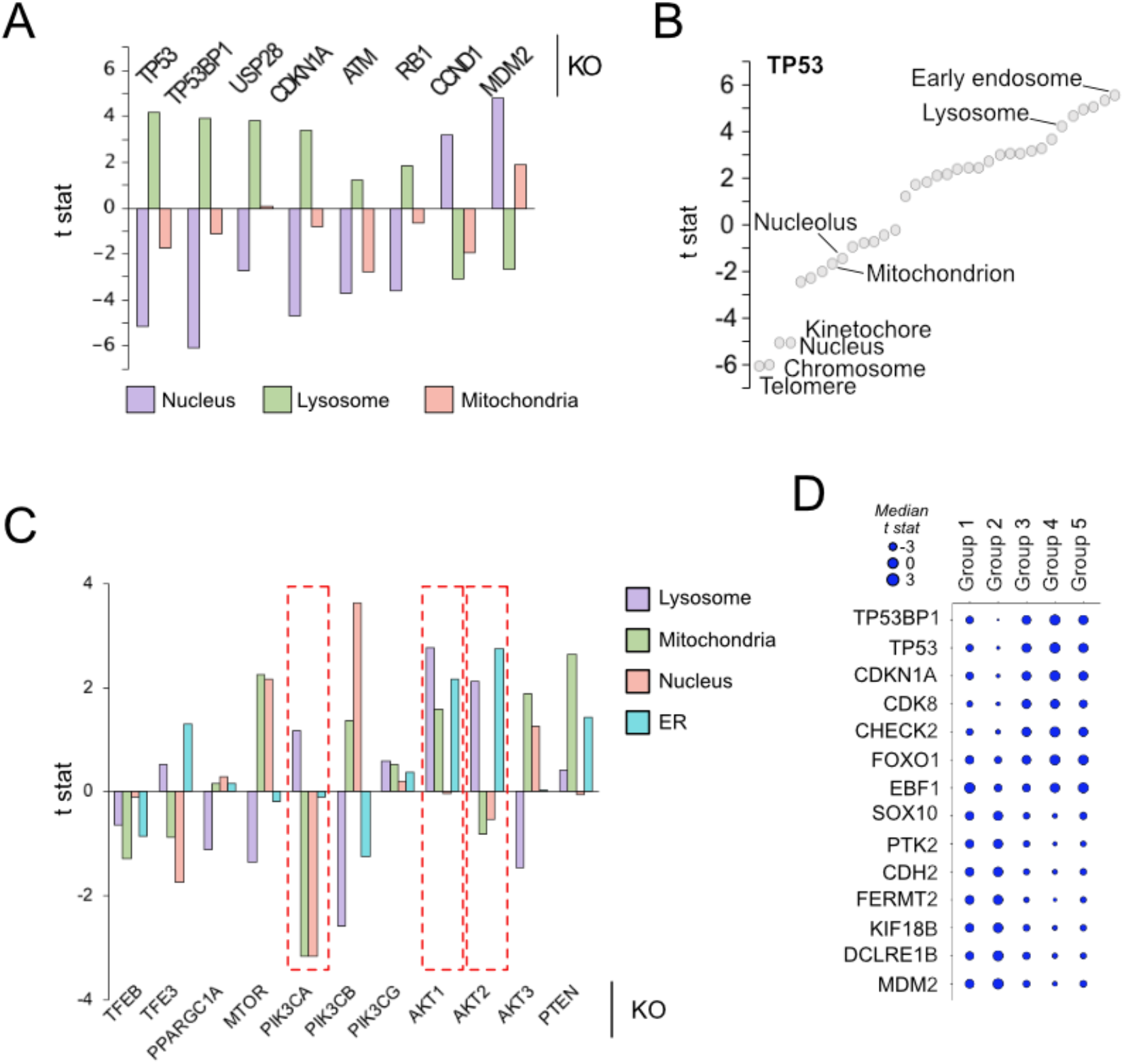
Organelle subproteome scaling dependencies. **(A)** Dependencies of nuclear, lysosomal and mitochondrial subproteomes on individual genes. **(B)** Dependencies of all subproteomes on TP53. **(C)** Dependencies of nuclear, lysosomal, mitochondrial and ER subproteomes on selected genes. Note the inverse dependencies on PI3K and AKT kinases (red dotted boxes). **(D)** Contributions of other genes into Organelle Scaling Dependency Map. Groups are the same as illustrated in the main fig. 4B.

## References

1. M. D’Ario et al., Cell size controlled in plants using DNA content as an internal scale. Science 372, 1176–1181 (2021).

2. K. M. Schmoller, J. J. Turner, M. Koivomagi, J. M. Skotheim, Dilution of the cell cycle inhibitor Whi5 controls budding-yeast cell size. Nature 526, 268–272 (2015).

3. E. Zatulovskiy, S. Zhang, D. F. Berenson, B. R. Topacio, J. M. Skotheim, Cell growth dilutes the cell cycle inhibitor Rb to trigger cell division. Science 369, 466–471 (2020).

4. W. F. Marshall, Scaling of Subcellular Structures. Annu Rev Cell Dev Biol 36, 219–236 (2020).

5. Y. H. Chan, W. F. Marshall, Scaling properties of cell and organelle size. Organogenesis 6, 88–96 (2010).

6. T. P. Miettinen, M. J. Caldez, P. Kaldis, M. Bjorklund, Cell size control - a mechanism for maintaining fitness and function. Bioessays 39, (2017).

7. Y. Gu, S. Oliferenko, The principles of cellular geometry scaling. Curr Opin Cell Biol 68, 20–27 (2020).

8. S. Kaushik, A. M. Cuervo, Proteostasis and aging. Nat Med 21, 1406–1415 (2015).

9. D. McHugh, J. Gil, Senescence and aging: Causes, consequences, and therapeutic avenues. J Cell Biol 217, 65–77 (2018).

10. A. Hernandez-Segura, J. Nehme, M. Demaria, Hallmarks of Cellular Senescence. Trends Cell Biol 28, 436–453 (2018).

11. N. E. Sharpless, C. J. Sherr, Forging a signature of in vivo senescence. Nat Rev Cancer 15, 397–408 (2015).

12. G. E. Neurohr et al., Excessive Cell Growth Causes Cytoplasm Dilution And Contributes to Senescence. Cell 176, 1083–1097 e1018 (2019).

13. H. An, A. Ordureau, M. Korner, J. A. Paulo, J. W. Harper, Systematic quantitative analysis of ribosome inventory during nutrient stress. Nature 583, 303–309 (2020).

14. S. C. Dolfi et al., The metabolic demands of cancer cells are coupled to their size and protein synthesis rates. Cancer Metab 1, 20 (2013).

15. T. Guo et al., Quantitative Proteome Landscape of the NCI-60 Cancer Cell Lines. iScience 21, 664–680 (2019).

16. M. C. Lanz et al., Increasing cell size remodels the proteome and promotes senescence. BioRxiv preprint https://doi.org/10.1101/2021.07.29.454227, (2021).

17. S. Rodriguez-Lopez et al., Mitochondrial adaptations in liver and skeletal muscle to prolongevity nutritional and genetic interventions: the crosstalk between calorie restriction and CYB5R3 overexpression in transgenic mice. Geroscience 42, 977–994 (2020).

18. R. Kafri et al., Dynamics extracted from fixed cells reveal feedback linking cell growth to cell cycle. Nature 494, 480–483 (2013).

19. D. F. Berenson, E. Zatulovskiy, S. Xie, J. M. Skotheim, Constitutive expression of a fluorescent protein reports the size of live human cells. Mol Biol Cell 30, 2985–2995 (2019).

20. P. Jorgensen et al., The size of the nucleus increases as yeast cells grow. Mol Biol Cell 18, 3523–3532 (2007).

21. C. D. Wiley et al., Analysis of individual cells identifies cell-to-cell variability following induction of cellular senescence. Aging Cell 16, 1043–1050 (2017).

22. R. Tacutu et al., Human Ageing Genomic Resources: new and updated databases. Nucleic Acids Res 46, D1083–D1090 (2018).

23. J. Lahtela et al., A high-content cellular senescence screen identifies candidate tumor suppressors, including EPHA3. Cell Cycle 12, 625–634 (2013).

24. J. C. Acosta et al., A complex secretory program orchestrated by the inflammasome controls paracrine senescence. Nat Cell Biol 15, 978–990 (2013).

25. T. P. Miettinen, M. Bjorklund, Cellular Allometry of Mitochondrial Functionality Establishes the Optimal Cell Size. Dev Cell 39, 370–382 (2016).

26. T. P. Miettinen, M. Bjorklund, Mitochondrial Function and Cell Size: An Allometric Relationship. Trends Cell Biol 27, 393–402 (2017).

27. A. Y. Chang, W. F. Marshall, Dynamics of living cells in a cytomorphological state space. Proc Natl Acad Sci U S A 116, 21556–21562 (2019).

28. D. L. Levy, R. Heald, Nuclear size is regulated by importin alpha and Ntf2 in Xenopus. Cell 143, 288–298 (2010).

29. K. A. Hansson et al., Myonuclear content regulates cell size with similar scaling properties in mice and humans. Nat Commun 11, 6288 (2020).

30. Y. H. Chan, W. F. Marshall, Organelle size scaling of the budding yeast vacuole is tuned by membrane trafficking rates. Biophys J 106, 1986–1996 (2014).

31. E. M. Jack et al., Increase in hepatocyte and nuclear volume and decrease in the population of binucleated cells in preneoplastic foci of rat liver: a stereological study using the nucleator method. Hepatology 11, 286–297 (1990).

32. E. M. Jack et al., Ultrastructural changes in chemically induced preneoplastic focal lesions in the rat liver: a stereological study. Carcinogenesis 11, 1531–1538 (1990).

33. D. Larrieu, S. Britton, M. Demir, R. Rodriguez, S. P. Jackson, Chemical inhibition of NAT10 corrects defects of laminopathic cells. Science 344, 527–532 (2014).

34. S. Bernales, K. L. McDonald, P. Walter, Autophagy counterbalances endoplasmic reticulum expansion during the unfolded protein response. PLoS Biol 4, e423 (2006).

35. A. Khaminets et al., Regulation of endoplasmic reticulum turnover by selective autophagy. Nature 522, 354–358 (2015).

36. F. Fumagalli et al., Translocon component Sec62 acts in endoplasmic reticulum turnover during stress recovery. Nat Cell Biol 18, 1173–1184 (2016).

37. X. Jiang et al., FAM134B oligomerization drives endoplasmic reticulum membrane scission for ER-phagy. EMBO J 39, e102608 (2020).

38. E. R. Weibel, C. R. Taylor, H. Hoppeler, The concept of symmorphosis: a testable hypothesis of structure-function relationship. Proc Natl Acad Sci U S A 88, 10357–10361 (1991).

39. M. P. Viana et al., Robust integrated intracellular organization of the human iPS cell: where, how much, and how variable? biorxiv, (2020).

40. M. Ghandi et al., Next-generation characterization of the Cancer Cell Line Encyclopedia. Nature 569, 503–508 (2019).

41. D. P. Nusinow et al., Quantitative Proteomics of the Cancer Cell Line Encyclopedia. Cell 180, 387–402 e316 (2020).

42. A. E. Mazzucco et al., Genetic interrogation of replicative senescence uncovers a dual role for USP28 in coordinating the p53 and GATA4 branches of the senescence program. Genes Dev 31, 1933–1938 (2017).

43. Z. Wu et al., Mechanisms controlling mitochondrial biogenesis and respiration through the thermogenic coactivator PGC-1. Cell 98, 115–124 (1999).

44. M. Sardiello et al., A gene network regulating lysosomal biogenesis and function. Science 325, 473–477 (2009).

45. K.-L. Claude, D. Bureik, P. Adarska, A. Singh, K. M. Schmoller, Transcription coordinates histone amounts and genome content. BioRxiv preprint https://doi.org/10.1101/2020.08.28.272492, (2021).

## References

1. S. C. Dolfi et al., The metabolic demands of cancer cells are coupled to their size and protein synthesis rates. Cancer Metab 1, 20 (2013).

2. T. Guo et al., Quantitative Proteome Landscape of the NCI-60 Cancer Cell Lines. iScience 21, 664–680 (2019).

3. D. C. Zielinski et al., Systems biology analysis of drivers underlying hallmarks of cancer cell metabolism. Sci Rep 7, 41241 (2017).

4. E. Eden, R. Navon, I. Steinfeld, D. Lipson, Z. Yakhini, GOrilla: a tool for discovery and visualization of enriched GO terms in ranked gene lists. BMC Bioinformatics 10, 48 (2009).

5. D. Szklarczyk et al., STRING v11: protein-protein association networks with increased coverage, supporting functional discovery in genome-wide experimental datasets. Nucleic Acids Res 47, D607–D613 (2019).

6. B. DepMap, DepMap 20Q2 Public. figshare. Dataset. https://doi.org/10.6084/m9.figshare.12280541.v4., (2020).

7. M. Ghandi et al., Next-generation characterization of the Cancer Cell Line Encyclopedia. Nature 569, 503–508 (2019).

8. D. P. Nusinow et al., Quantitative Proteomics of the Cancer Cell Line Encyclopedia. Cell 180, 387–402 e316 (2020).

9. T. P. Miettinen, M. Bjorklund, Cellular Allometry of Mitochondrial Functionality Establishes the Optimal Cell Size. Dev Cell 39, 370–382 (2016).

10. E. M. Jack et al., Increase in hepatocyte and nuclear volume and decrease in the population of binucleated cells in preneoplastic foci of rat liver: a stereological study using the nucleator method. Hepatology 11, 286–297 (1990).

11. E. M. Jack et al., Ultrastructural changes in chemically induced preneoplastic focal lesions in the rat liver: a stereological study. Carcinogenesis 11, 1531–1538 (1990).

